# Transient Dietary Changes Modulate Inflammatory Disease Trajectory in the Lean State

**DOI:** 10.1101/2025.06.15.659784

**Authors:** Zewen Jiang, John M. Tayeri, Chihiro Tabuchi, Sarah G. Gayer, Arum Yoo, Im-Hong Sun, Jiaxi Wang, Han Yin, Shin Shyan Chuah, Preetal Deshpande, Vrinda Johri, Brittany Davidson, Jessica Tsui, Nicholas F. Kuhn, Tristan Courau, Alexis J. Combes, James M. Gardner, Sagar P. Bapat

**Affiliations:** Diabetes Center, University of California, San Francisco, CA, USA; Department of Laboratory Medicine, University of California, San Francisco, CA, USA; Department of Pathology, University of California, San Francisco, CA, USA; ImmunoX Initiative, University of California, San Francisco, CA, USA; UCSF CoLabs, University of California, San Francisco, CA, USA; Department of Medicine, University of California, San Francisco, CA, USA; Department of Surgery, University of California, San Francisco, CA, USA

## Abstract

Obesity potently alters immune responses across various inflammatory contexts^1–7^. However, it is unclear whether a transient high-fat diet (HFD) can affect immune function despite minimal effects on body weight. Here, we demonstrate that a short-term HFD regimen significantly exacerbates disease severity in a model of experimental psoriasis, comparable to what is observed in obese mice on long-term HFD. We find that a critical 4-day window of HFD coinciding with disease onset is sufficient to overactivate the immune response, leading to worsened disease outcomes. Mechanistically, we identify the CD4^+^ T cell compartment as an essential mediator of HFD-induced disease severity. Within the compartment, we functionally validate that disease exacerbation is driven by increased differentiation of a pathogenic population of T helper 17 (T_H_17) cells^8^ expressing IL1R1 (interleukin-1 receptor, type I)^9^, which is triggered by localized activation of the NLRP3 (NOD-, LRR-, and pyrin domain-containing protein 3) inflammasome^10^. This brief window of HFD at disease onset leads to immune-dependent sensitization, as disease severity is increased upon a subsequent flare, despite complete recovery and under a low-fat diet (LFD) regimen. Our data indicate that transient dietary changes during the initial stages of an inflammatory event can rewire the immune milieu, profoundly influencing disease progression and inflammatory memory^11^. This phenomenon may have broad implications for conditions where obesity, and not diet itself, is considered the primary risk factor.

## Main

Obesity substantially affects immune function across diverse inflammatory contexts, including infection, autoimmunity, and cancer^7,12–17^. This is particularly evident in cutaneous inflammation, where obesity can exacerbate disease severity and complicate therapeutic response^6,18^. While more is known about the inflammatory impact of obesity, the effects of transient, high-fat dietary shifts on immune regulation and disease progression, in the lean state, remain poorly understood. Psoriasis represents an ideal model to examine these short-term dietary influences, as it is both intimately linked with obesity^4,15^ and known to flare in response to poorly defined nutritional triggers^19–22^. Here, we ask whether a brief bout of high-fat feeding, which is not sufficient to cause significant weight gain, can reshape immune responses in psoriatic inflammation and potentially reprogram the long-term trajectory of this disease.

### Transient HFD is comparable to long-term HFD in exacerbating psoriasis

To differentiate the effects of a short-term diet from those of obesity on inflammatory responses, we employed an experimental model of psoriasis. Age-matched mice were subjected to either a short-term (1-week) or long-term (10-week) HFD (versus mice fed LFD as controls), followed by the induction of psoriasis-like inflammation via daily topical imiquimod (IMQ) application to the ear (**Fig. 1a**). Notably, the 1-week HFD regimen did not result in significant weight gain compared to the LFD regimen, whereas the 10-week HFD induced marked obesity (**Fig. 1b**). Surprisingly, both dietary regimens resulted in a similar, significant (∼2.1-fold for 1-week HFD over LFD and ∼2.3 fold for 10-week HFD over LFD) increase in change in ear thickness relative to LFD-fed mice (**Fig. 1c**). Macroscopic examination showed a reduction in ear length from base to tip, and histological analysis of ear skin cross-sections revealed substantial expansion of both the dermis and epidermis under both dietary regimens (**Fig. 1d**). Since IMQ-induced psoriatic inflammation predominantly stimulates Type 3 immune responses, characterized by the production of IL-17A, IL-17F, and IL-22 by T_H_17 cells, γδ T cells, and other immune cells^23,24^, we used flow cytometry to quantify these populations (gating strategy shown in **Extended Data Fig. 1**). Both 1-week and 10-week HFD feeding resulted in significantly elevated levels of T_conv_ (FOXP3^−^ CD4^+^) cells expressing the lineage defining transcription factor for T_H_17 cells RORγt and able to produce the T_H_17 cellular effector cytokines IL-17A, IL-17F, and IL-22 in lesional skin (**Fig. 1e,f**) and in skin draining lymph nodes (**Extended Data Fig. 2a,b**). The 1-week HFD regimen significantly increased the proliferative capacity of CD4⁺ T cells, as indicated by Ki-67 expression, to levels comparable to those in obese mice (**Fig. 1g**). Similarly, the proportion of regulatory T (T_reg_) cells among CD4⁺ T cells were decreased in both short-term and long-term HFD-fed mice (**Fig. 1h**). Neutrophils, key mediators of Type 3 inflammation^25,26^, were also significantly elevated after just 1 week of HFD, mirroring levels observed in long-term HFD-fed obese mice (**Fig. 1i**). By contrast, the frequency of γδ T cells that were competent for IL-17A, IL-17F, and IL-22 did not increase upon HFD feeding (**Extended Data Fig. 2c**), although the total number of γδ T cells in the skin of both groups increased, relative to their LFD-fed controls (**Extended Data Fig. 2d**). In summary, these findings demonstrate that short-term HFD exposure can intensify psoriatic inflammation to levels comparable to long-term HFD-induced obesity, suggesting that obesogenic diets directly influence immune responses independently of obesity.

**Figure 1.**
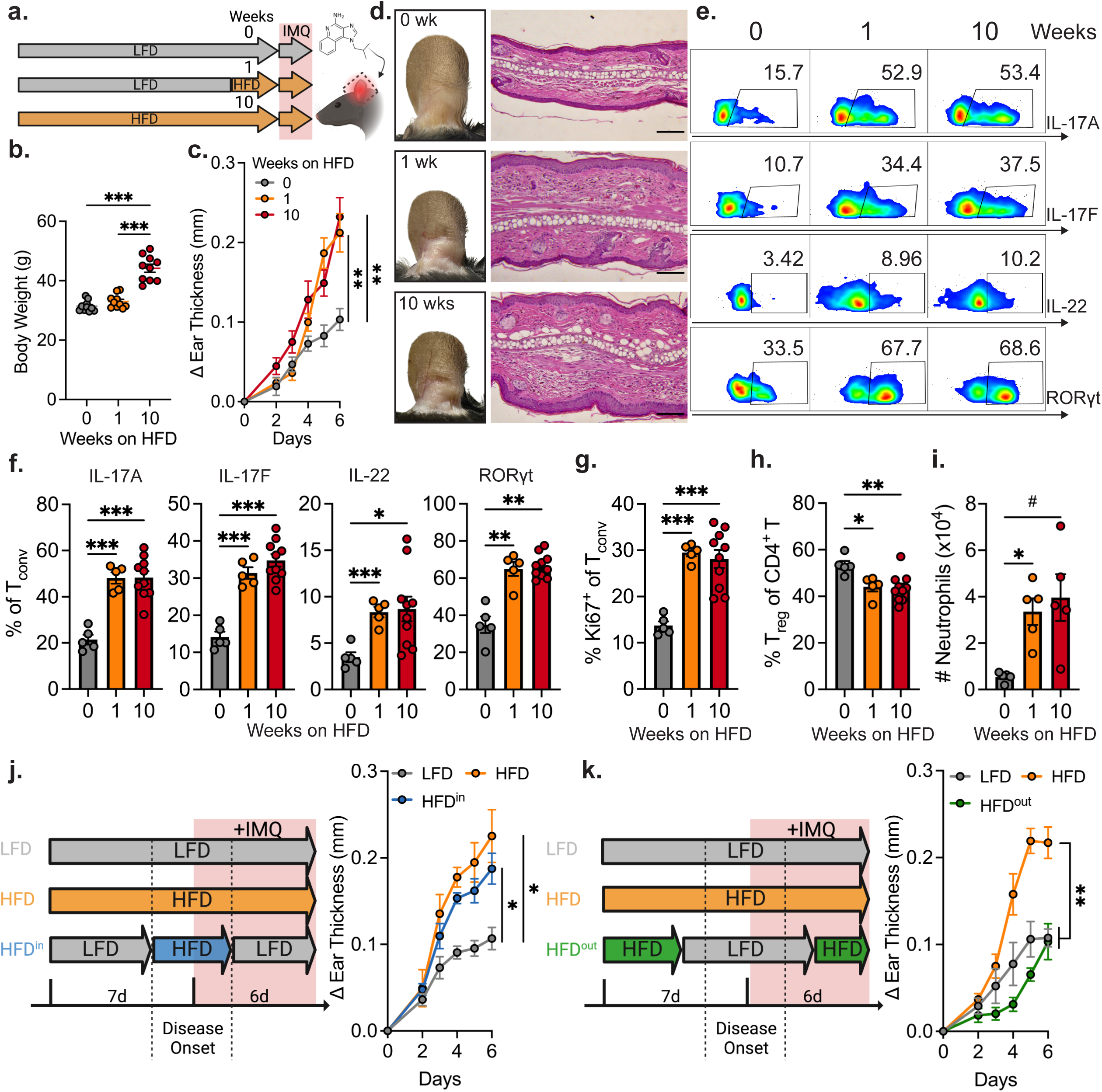
Transient, short-term HFD intensifies psoriatic inflammation comparable to obesity, **a**, Imiquimod (IMQ)-induced psoriasis model with either low-fat diet (LFD) or high-fat diet (HFD) feeding for indicated weeks, **b**, Body weight after LFD or HFD feeding for indicated weeks before IMQ challenge, **c**, Change in ear thickness during disease development, **d**, Representative ear images and hematoxylin and eosin (H&E) ear histology of mice on day 6 of IMQ treatment, **e**, Representative flow cytometry plots of IL-17A^+^, IL-17F^+^, IL-22^+^, and RORγt^+^ conventional T (T_conv_) cells (defined as CD4^+^ FOXP3) in the skin, **f**, Quantification of (**e**) by percentage, **g**. Percentage of Ki-67^+^ T_conv_ cells, **h**, Percentage of FOXP3^+^ T_reg_ cells (out of CD4^+^ T cells) in the skin, **i**, Total number of neutrophils (defined as CD45^+^ CD3ε^−^ CD11b^+^ Ly6G^+^) in the skin, **j**, Schematic of IMQ-induced psoriasis with HFD feeding within disease onset (HFD^in^) and change in ear thickness during disease development with HFD^in^ feeding. **k**, Schematic of IMQ-induced psoriasis with HFD feeding outside of disease onset (HFD^out^) and change in ear thickness during disease development with HFD^out^ feeding. Scale bars, 100 µm. n=10 **(b,c**); n=5 for 0 and 1 weeks groups and n=10 for 10 weeks group (**f-h**); n=5 (**i-k**). Data are mean ± s.e.m. Brown-Forsythe and Welch ANOVA test was performed followed by post-hoc pairwise comparisons using Dunnett’s T3 multiple comparison test to calculate P values (**b,c,f-k**). Only peak values were tested in (**cj,k**). *^#^P* = 0.07, *\*P* < 0.05, *\*\*P* < 0.01, *\*\*\*P* < 0.001.

**Extended Data Figure 1.**
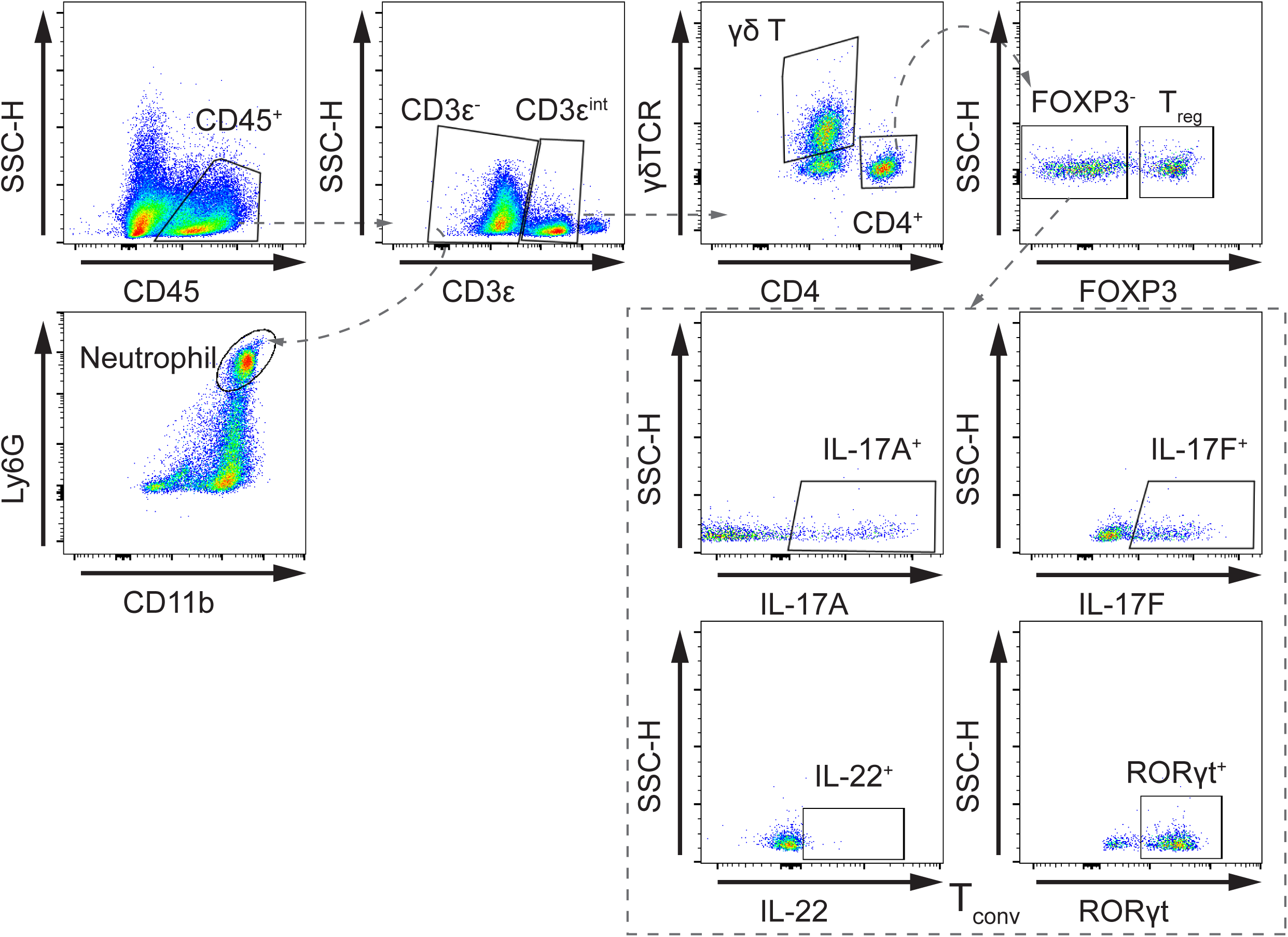
Selected flow cytometry gating strategies to quantify γδT cells, T_H_17 cells, T_reg_ cells, and neutrophils. Specifically conventional T cells (T_conv_) are defined as CD45^+^ CD3ε^lnt^CD4^+^ FOXP3^−^, T_H_17 cells are defined as T_conv_ cells expressing RORγt and competent for IL-17A, IL-17F, and IL-22, γδ T cells are defined as CD45^+^ CD3ε^lnt^γδTCR^+^, and neutrophils are defined as CD45^+^ CD3ε^−^ CD11 b^+^ Ly6G^+^.

**Extended Data Figure 2.**
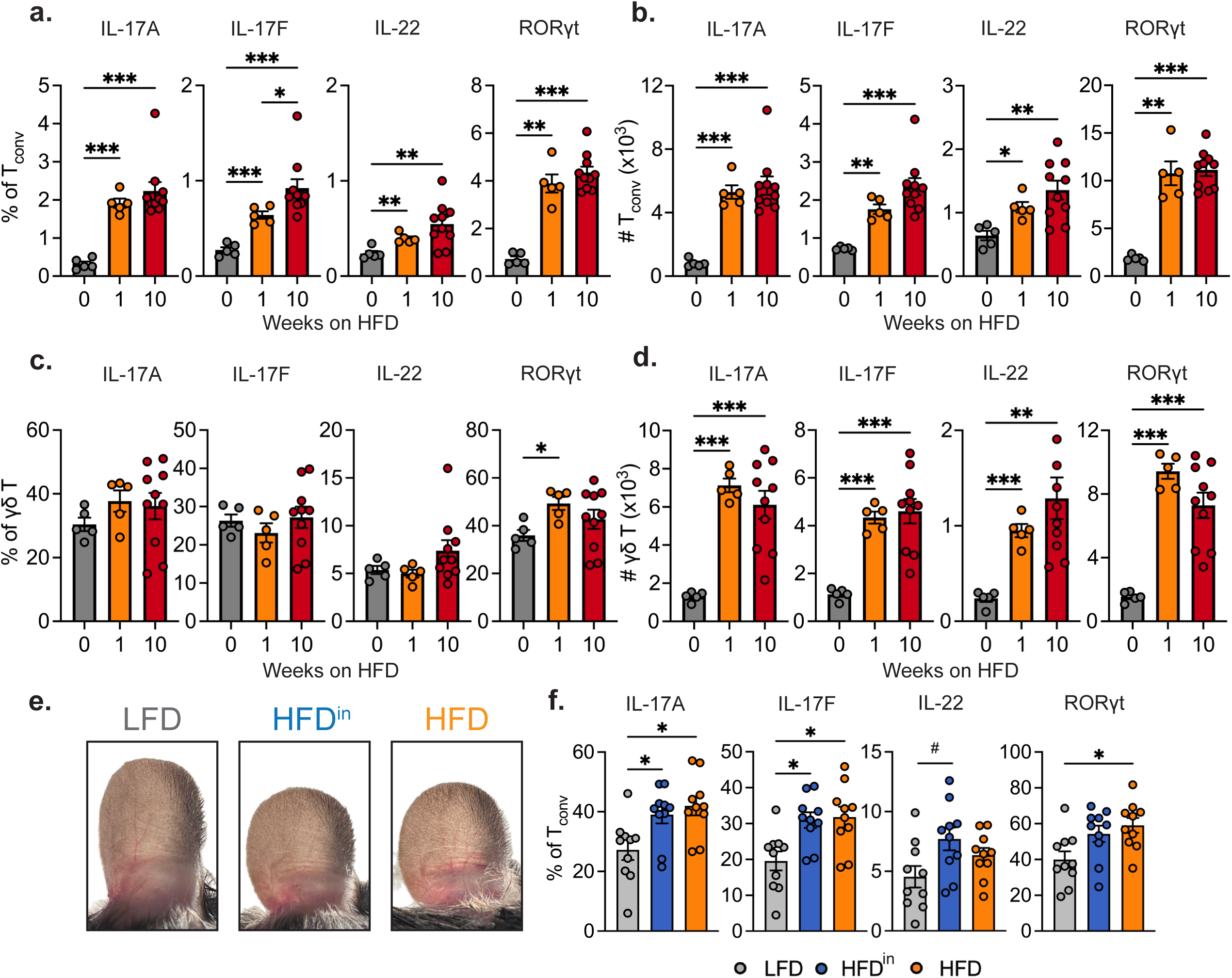
Transient, short-term HFD increases Type 3 inflammation upon IMQ treatment, **a,b,** Percentage **(a)** and total number **(b)** of IL-17A^+^, IL-17F^+^, IL-22^+^, and RORγt^+^ T_conv_ cells (defined as CD4^+^ FOXP3^−^) in the skin draining lymph nodes (n=5 for 0,1 weeks and n=1Ū for 10 weeks), **c,d,** Percentage **(c)** and total number **(d)** of IL-17A^+^, IL-17F^+^, IL-22^+^, and RORγt^+^γδ T cells in the skin (n=5 for 0,1 weeks and n=10 for 10 weeks), e, Representative ear images of mice fed either LFD, HFD^in^, or HFD on day 6 of IMQ treatment, **f,** Percentage of IL-17A^+^, IL-17F^+^, IL-22^+^, and RORγt^+^ T_conv_ in the skin, n=5 for 0,1 weeks and n=10 for 10 weeks **(a-d);** n=10 **(f).** Data are mean ± s.e.m. Brown-Forsythe and Welch ANOVA test was performed followed by post-hoc pairwise comparisons using Dunnett’s T3 multiple comparison test to calculate P values **(a-d,f).** *^#^P =* 0.07, *\*P<* 0.05, **P< 0.01, *\*\*\*p<* 0.001.

### HFD at disease onset is sufficient to drive exacerbated skin inflammation

To examine the dynamics of this short-term HFD-induced Type 3 inflammation, we further investigated the temporal relationship between HFD-feeding and IMQ-induced inflammatory disease severity. Remarkably, 4 days of HFD feeding around the initiation of disease (HFD^in^) were sufficient to cause a significant escalation in psoriatic inflammation (**Fig. 1j, Extended Data Fig. 2e,f**). By contrast, HFD-feeding given outside this critical window (HFD^out^) did not significantly impact disease severity compared to mice maintained on LFD throughout (**Fig. 1k**). Overall, these results suggest that diet, specifically diet coincident with the initiation of an inflammatory cascade, can potently alter inflammatory disease trajectory.

### CD4^+^ T cells are required for disease exacerbation following short-term HFD

Next, we sought to identify the critical immune cell types that enhance disease severity and Type 3 inflammation upon HFD-feeding. To this end, we first used *Rag1*^⁻/⁻^ mice, which lack adaptive immunity but retain innate lymphoid cells (ILCs)^27^. These mice were resistant to HFD-induced exacerbation of disease severity, suggesting that T or B cells, but not ILCs, are required for this effect (**Fig. 2a,b**). Given the known role of γδ T cells as major IL-17 producers in the skin and their significant expansion upon long-term HFD feeding^18,28–31^, we examined the effects of short-term HFD on *Tcrd*^−/−^ mice, which do not develop γδ T cells, upon IMQ challenge. Surprisingly, HFD-fed *Tcrd*^−/−^ mice developed significantly greater disease compared to LFD-fed controls (**Fig. 2c**), suggesting that γδ T cells are dispensable for the short-term HFD-induced inflammatory exacerbation. We next investigated the role of αβ T cells in this process using *Tcra*^−/−^ mice, which do not develop αβ T cells, and did not observe differences between mice fed HFD versus LFD. To further identify the subset within the αβ T cell compartment that is required for the HFD-enhancement of disease severity, we utilized *B2m*^−/−^ mice, which are deficient in CD8^+^ T cells, and MHCII^Δ/Δ^ mice, which lack CD4^+^ T cells. Notably, only CD4^+^ T cell deficiency resulted in the loss of the exacerbated disease and inflammatory response caused by HFD in wild-type (WT) mice (**Fig. 2d-f**). To further confirm the essential role of CD4^+^ T cells in altering disease trajectory in response to HFD, we treated WT mice with IgG or anti-CD4 antibodies. Anti-CD4 antibody treatment completely abolished the diet-induced difference in disease severity between mice fed LFD versus HFD (**Extended Data Fig. 3a-d**). We then sought to identify whether the presence of CD4^+^ T cells, in the absence of other adaptive immune cells, could produce an exaggerated immune response to HFD. We adoptively transferred bulk CD4^+^ T cells isolated from the spleen of WT mice into *Rag1*^−/−^ mice and, upon homeostatic expansion, provided HFD or LFD for 1 week, followed by IMQ treatment (**Fig. 2g**). *Rag1*^−/−^ mice that were transferred with CD4^+^ T cells had a robust enhancement of IMQ-induced disease upon HFD feeding with elevated numbers of T_H_17 cells in the skin (as marked by RORγt-positivity and FOXP3-negativity) (**Fig. 2h-k**). Taken together, these results demonstrate that CD4^+^ T cells are critical immunological mediators translating dietary influences into enhanced inflammatory responses in this model of inflammatory skin disease.

**Figure 2.**
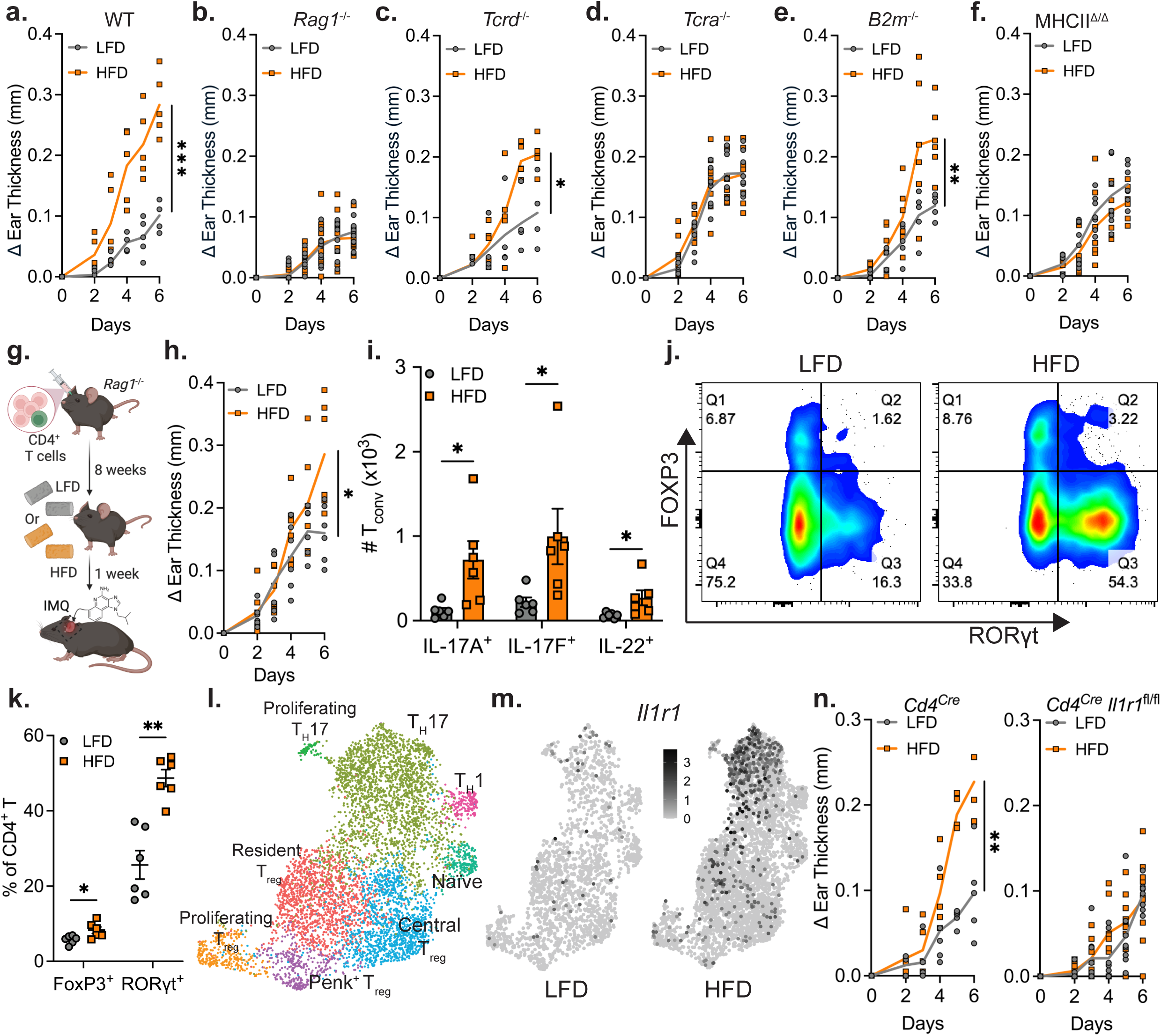
The exaggerated disease severity due to short-term HFD feeding is driven by IL1R1^+^ T_H_17 T cells, **a-f,** Change in ear thickness during disease development in WT mice **(a),** *Ragl^−/−^* mice **(b),** *Tcrd^−/−^* mice **(c),** *Tcra^−/−^* mice **(d),** *B2m^−/−^* mice **(e),** and MHCII^Δ/Δ^ mice **(f)** fed LFD or HFD. **g,** Schematic of bulk CD4^+^ T cell adoptive transfer, **h,** Change in ear thickness over time in adoptive transfer recipient mice fed LFD or HFD. **i,** Total number of IL-17A^+^, IL-17F^+^, and IL-22^+^ T_conv_ cells in the skin, **j,** Representative flow cytometry plots of FOXP3 and RORγt expression in CD4^+^ T cells in the skin, k, Quantification of (j) by percentage. I, Visualization in UMAP space with Leiden clustering of scRNA-seq data of CD4^+^ T cells from the skin of LFD- or HFD-fed WT mice treated with IMQ for 6 days, **m,** Gene expression heatmap in UMAP space (from I) of *IHr1* with the highest-expressing cells in black, n, Change in ear thickness during disease development in *Cd4^cre^* mice or IL1 R1-deficient *(Cd4^cre^ IHr1™)* mice fed LFD or HFD. *n=5* **(a);** n=10 **(b);** *n=4* for LFD and n=5 for HFD **(c);** n=8 **(d);** n=5 for LFD and n=6 for HFD **(e);** n=7 **(f);** n=6 **(h,i,k);** n=1 (pooled from 10 mice per group) for scRNA-seq **(l,m);** n=5 for *Cd4^cre^, n=8* for LFD *Cd4^cre^ IHr1™,* and n=9 for HFD *Cd4^cre^ IHr1™* (n). Data are mean ± s.e.m. Welch’s ŕ-test was used to calculate *P* values, except the Mann-Whitney test was used for **(i).** P values were corrected for multiple corrections using Holm-Šídák method **(i,k).** Only peak values were tested in **(a-f,h,n).** *\*P <* 0.05, *\*\*P* < 0.01, ***P < 0.001.

**Extended Data Figure 3.**
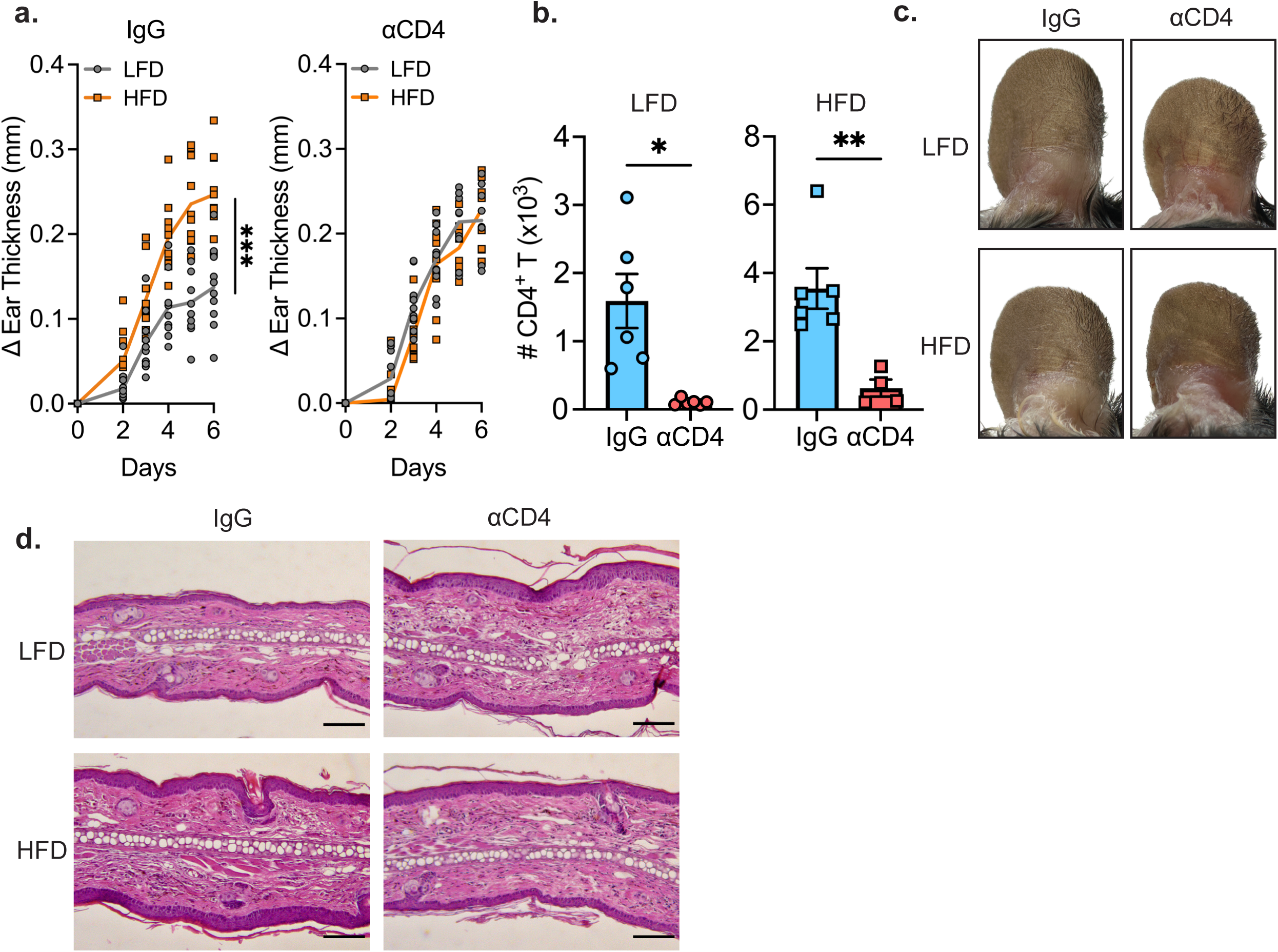
Treatment with neutralizing anti-CD4 antibody eliminates diet-associated inflammatory differences, **a,b,** Change in ear thickness **(a)** and total number of lesional CD4^+^ T cells **(b)** in mice treated with IgG isotype control or anti-CD4 antibody and challenged with IMQ. **c**, Representative ear images of mice onday 6 of IMQ treatment upon either IgG or anti-CD4 antibody treatment, **d,** Representative hematoxylin and eosin (H&E) ear histology of mice on day 6 of IMQ treatment upon either IgG or anti-CD4 antibody treatment. Scale bars, 100 µm. n=10 (a); n=6 for LFD IgG, LFD anti-CD4 antibody, and HFD IgG treatment and n=4 for HFD anti-CD4 antibody treatment **(b).** Data are mean ± s.e.m. Welch’s ŕ-test was used to calculate P values. Only peak values were tested in **(a).** *P* values were corrected for multiple corrections using Holm-Šídák method (b). *P < 0.05, *\*\*P* < 0.01, *\*\*\*P* < 0.001.

### IL1R1^+^ CD4^+^ T cells transduce the exaggerated inflammatory effects of short-term HFD

To further elucidate the mechanistic underpinnings of how CD4^+^ T cells are influenced by short-term HFD feeding, we systematically profiled the lesional CD4^+^ T cells from LFD and HFD-fed mice after IMQ-challenge using single-cell RNA sequencing (scRNA-seq). Exploratory analysis revealed 3 T_conv_ cell clusters (T_H_1, T_H_17, and proliferating T_H_17), 4 distinct T_reg_ cell clusters (Proliferating, Resident, Central, and Penk^+32^), and a naïve cluster (**Fig. 2l**). The gene-expression heatmaps of effector cytokines (*Il17a*, *Il17f*, *Il22*, *Ifng*, and *Fasl)*^33,34^, key transcription factors (*Rorc*, *Tbx21, Foxp3*, and *Ikzf2*)^35–37^, cytokine receptors (*Il23r*, *Il1rl1*, *Il1r1*, *and Il2ra*)^31,38^, markers of proliferation (*Mki67*, *Cdca3*, *Spc24*, and *Pclaf*), prominent markers of activation, quiescence, tissue residency, and memory (*Ccr7*, *Sell*, *Itgae*, *Klf2*, and *Bcl2*)^6^, and *Penk*^32^ validated these population assignments (**Extended Data Fig. 4a-f**). Cells from mice fed HFD showed marked differences in distribution across the cell populations, particularly notable in the T_H_17 cluster (**Extended Data Fig. 5a**). In cells that have T_H_17 identities (including T_H_17 and proliferating T_H_17), T_H_17 effector genes, such as *Il17a*, *Il17f*, *Il22*, *Il23r* and *Rorc*, were significantly upregulated in CD4^+^ T cells from HFD-fed mice (**Extended Data Fig. 5b**). Furthermore, leveraging published gene signatures associated with T_H_17 pathogenicity^8,39^, we found that short-term HFD feeding markedly enhances pathogenicity of T_H_17 cells in the skin (**Extended Data Fig. 5c**). We performed differential gene expression analysis (**Extended Data Fig. 5d**, **Supplementary Table 1**) and found that *Il1r1* was the most significantly upregulated gene from T_H_17 cells taken from mice fed HFD (**Fig. 2m, Extended Data Fig. 5d**). These findings were further validated by flow cytometry, which confirmed that HFD feeding significantly increases IL1R1 expression in CD4^+^ T cells from lesional skin (**Extended Data Fig. 5e**). IL1R1 signaling has been shown to be critical for stabilizing and enhancing Type 3 inflammation^9,40–42^. CD4^+^ T cells expressing *Il1r1* exhibited increased expression of T_H_17 effector genes, along with a higher T_H_17 pathogenicity score (**Extended Data Fig. 5f,g**). T_H_17 differentiation involves multiple inflammatory cues, whose circuitry is well-defined *in vitro* but less well-understood in *in vivo* contexts^43,44^. These findings suggest that IL1R1 plays a specific and critical role in driving T_H_17 differentiation in mice being fed HFD. To directly test this hypothesis, we subjected mice with a T cell-specific ablation of IL1R1 (*Cd4^Cre^ Il1r1^fl/fl^*) to 1 week of LFD or HFD feeding, followed by IMQ challenge. Strikingly, T cell-specific IL1R1-deficient mice were completely protected from HFD-induced exaggeration of psoriatic inflammation, maintaining disease severity levels similar to LFD-fed controls (**Fig. 2n**, **Extended Data Fig. 5h**). Collectively, these results demonstrate that short-term HFD exacerbates psoriatic-like inflammation via IL1R1^+^ T_H_17 cells in the skin lesion.

**Extended Data Figure 4.**
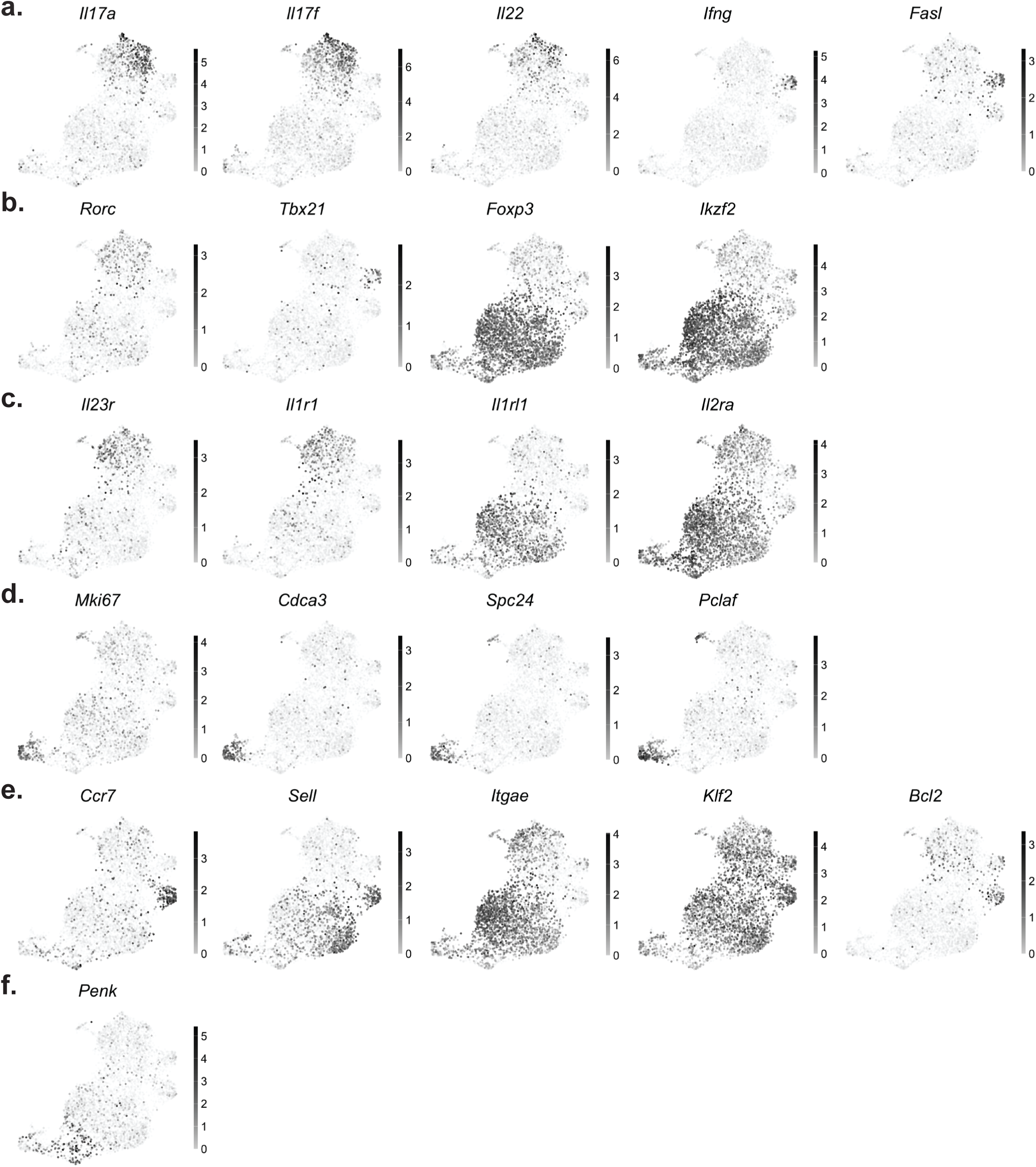
Gene heatmaps from scRNA-Seq data used in Fig. 2. **a-f**, Heatmaps of cytokines **(a),** transcription factors **(b),** cytokine receptors **(c),** markers of proliferation **(d),** markers of activation, quiescence, tissue residency, and memory **(e),** and *Penk* **(f)** overlain on UMAP plot from **Fig. 21** to assign names to clusters. Grayscale indicates gene expression, with the highest expressing cells in black.

**Extended Data Figure 5.**
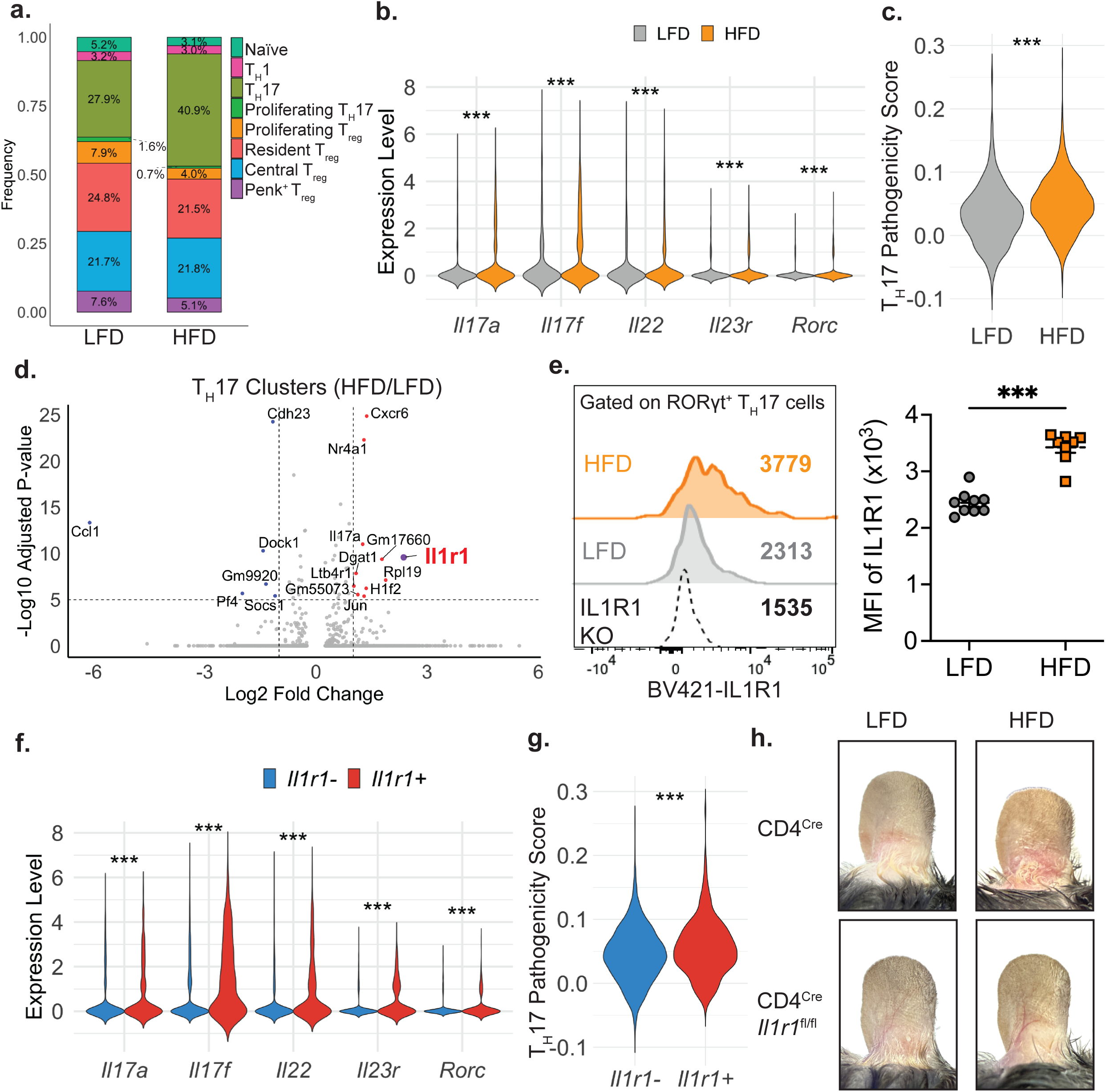
Short-term HFD feeding promotes psoriatic inflammation via IL1R1^+^ CD4^+^ T cells, **a-d,** Frequency of CD4^+^ T cells across assigned clusters **(a),** *II17a, II17f, II22, II23r,* and *Rorc* expression of CD4^+^ T cells from the T_H_17 clusters **(b),** T_H_17 pathogenicity score of CD4^+^ T cells in the T_H_17 clusters **(c),** and differential gene analysis in the T_H_17 clusters **(d)** as per clustering of scRNA-seq data in **Fig. 21. e,** Representative histogram of IL1R1 expression and quantification by mean fluorescence intensity (MFI) of RORγt^+^ Foxp3^−^ CD4^+^ T (T_H_17) cells from lesional skin, **f,g,** *II17*a*, II17f, II22, II23r,* and *Rorc* expression **(f)** and T_H_17 pathogenicity score **(g)** of *II1 r1^−^* and *II1r1^+^* T_H_17 cells as per clustering of scRNA-seq data in **Fig. 2I. h,** Representative ear images of *Cd4^cre^* or *Cd4^cre^ Iir1^fl/fl^* mice fed with either LFD or HFD on day 6 of IMQ treatment, *n-3* for LFD and *n-8* for HFD **(e).** Data are mean ± s.e.m. Welch’s f-test was used to calculate Rvalues for **(e)** and Wilcoxon test was used to calculate P values for **(b,c,f,g).** P values were corrected for multiple corrections using the Benjamini-Hochberg (BH) method **(b,f).** *\*P* < 0.05, *\*\*P* < 0.01, ***P < 0.001.

### Localized NLRP3-dependent inflammasome activation drives HFD-induced inflammatory exacerbation

We sought to identify the mechanistic basis by which IL1R1^+^ CD4^+^ T cells may be induced during IMQ challenge with short-term HFD-feeding. Since IL-1α and IL-1β are the known ligands for IL1R1^45^, we investigated whether HFD feeding combined with IMQ challenge increases the levels of these cytokines in the skin. We performed bulk RNA sequencing (RNA-seq) on whole ear tissues from mice fed either LFD or HFD, followed by IMQ challenge (**Supplementary Table 2**). In line with our earlier findings (**Fig. 1e,f, Extended Data Fig. 2a-d**), the ears of mice fed HFD exhibited a marked upregulation of Type 3 cytokine transcripts, including *Il17a* and *Il17f*, alongside genes involved in chemotaxis signaling such as *Cxcl2*, *Cxcl3*, *Ccl3*, and *Fpr1* (**Fig. 3a, Extended Data Fig. 6a**). Notably, *Il1b* exhibited dominant expression over *Il1a* in lesional ear skin, with HFD feeding inducing a robust ∼3.9 fold increase in *Il1b* expression (**Extended Data Fig. 6b**). Interestingly, this increase in *Il1b* was accompanied by a significant upregulation of *Nlrp3* (**Extended Data Fig. 6b**). Assembly of the canonical NLRP3-dependent inflammasome leads to the proteolytic cleavage of pro-caspase-1 into active caspase-1, which subsequently processes pro-IL-1β into its mature, active form^2,10^ (**Fig. 3b**) – leading us to hypothesize that the NLRP3-dependent inflammasome pathway might be important in driving differentiation of IL1R1^+^ T_H_17 cells in the skin of mice fed HFD during disease development. We observed a significant increase in both pro- and activated caspase-1 (p20) levels in the lesional skin-draining lymph nodes of HFD-fed mice, as compared to LFD-fed mice (**Fig. 3c**), but not from the contralateral skin-draining lymph nodes (**Extended Data Fig. 6c**) after IMQ challenge, suggesting HFD-feeding with IMQ challenge induces a localized activation of the NLRP3-dependent inflammasome. To directly test the importance of NLRP3 inflammasome activity in driving HFD-exaggerated disease, we subjected mice deficient in NLRP3 (*Nlrp3*^−/−^) to one week of either LFD or HFD feeding, followed by IMQ challenge. Remarkably, HFD-fed *Nlrp3*^−/−^ mice did not exhibit the exacerbated disease typically seen in WT HFD-fed mice, when compared to LFD-fed counterparts (**Fig. 3d-f**).

**Figure 3.**
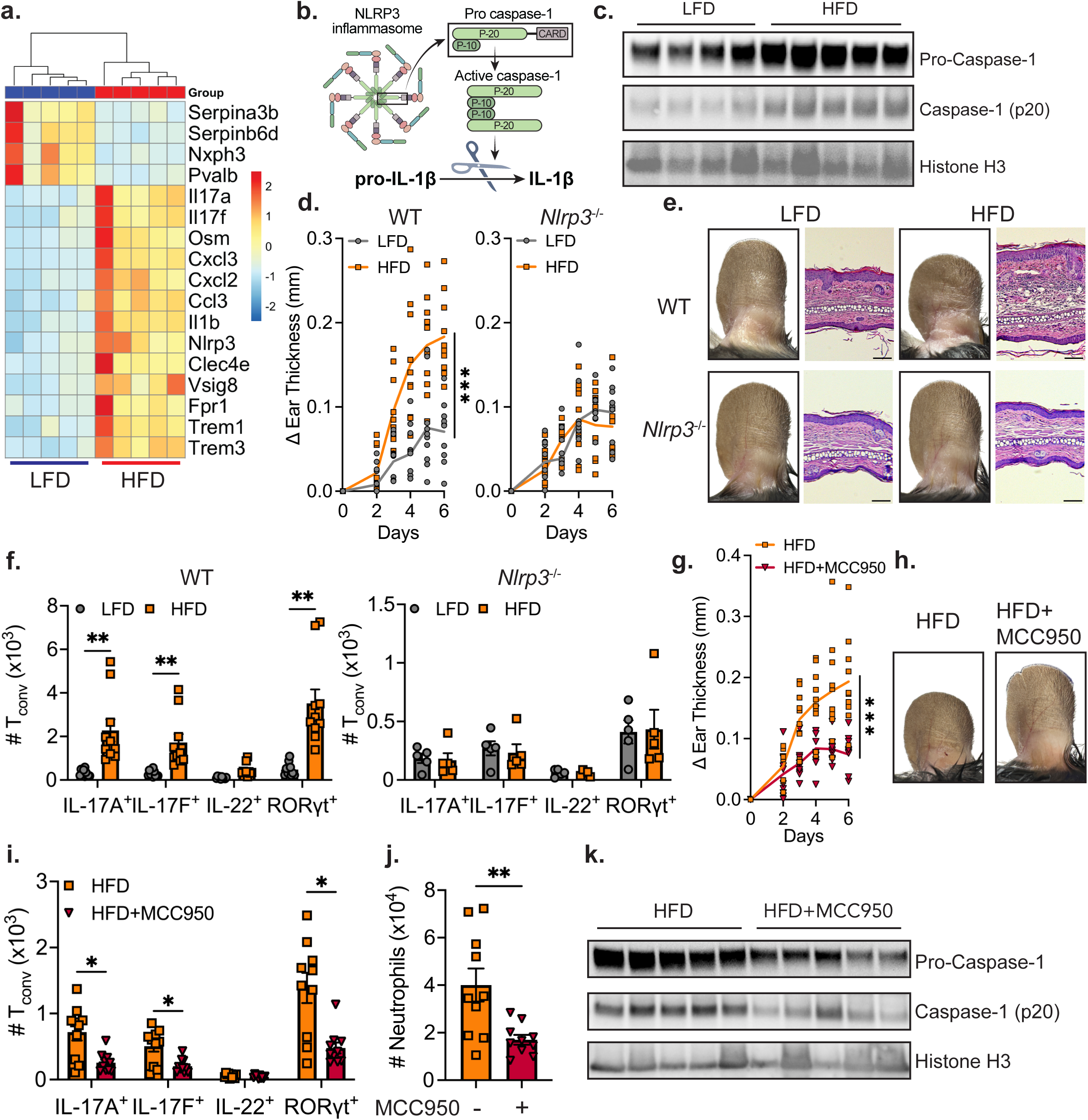
Transient HFD with IMQ challenge activates the NLRP3 inflammasome to drive the exaggerated inflammatory response, **a,** Expression heatmap of selected genes from RNA-seq of IMQ-treated ears of LFD or HFD-fed mice, **b,** Schematic of NLRP3 inflammasome activation, **c,** Western blot of Pro-Caspase-1, Capsase-1, and Histone H3 from skin draining lymph node lysates from WT mice fed LFD or HFD after treatment with IMQ. **d,** Change in ear thickness over time for WT or *Nlrp3’^1^’* mice fed LFD or HFD. **e,** Representative images and H&E histology of ears of WT or *Nlrp3^−/−^* mice on day 6 of IMQ treatment, f, Total number of IL-17A^+^, IL-17F^+^, IL-22^+^, and RORγt^+^ skin Tconv in WT or *Nlrp3^−/−^* mice, **g-k,** Change in ear thickness over time **(g),** representative ear images **(h),** total number of IL-17A^+^, IL-17F^+^, IL-22^+^, and RORγt^+^ skin T_conv_ (i) and skin neutrophils (j), and Western blot of Pro-Caspase-1, Capsase-1, and Histone H3 from skin draining lymph node lysates **(k)** of WT mice fed on HFD treated with vehicle or MCC950 with IMQ treatment. Scale bars, 100 µm. *n=5* (a,k); *n=4* for LFD, and *n=5* for HFD **(c);** n=10 **(d,f,g,i,j).** Data are mean ± s.e.m. Welch’s ŕ-test was used to calculate *P* values. *P* values were corrected for multiple corrections using Holm-Šídák method **(f,i).** Only peak values were tested in **(d,g).** *P < 0.05, *\*\*P* < 0.01, *\*\*\*P* < 0.001.

Although the NLRP3 inflammasome assembles and is active primarily in innate immune cells^10^, recent studies suggest a role for CD4^+^ T cell-intrinsic NLRP3-IL-1 signaling^46,47^. However, analysis of our scRNA-seq dataset of CD4^+^ T cells revealed negligible expression of *Nlrp3* in CD4⁺ T cells (**Extended Data Fig. 6d**). Consistent with this, both HFD-fed MHC II^Δ/Δ^ mice and, separately, HFD-fed *Cd4^Cre^ Il1r1^fl/fl^* mice retained intact caspase-1 processing, after IMQ challenge, suggesting that HFD-induced NLRP3 activation is extrinsic to and upstream of the involvement of CD4⁺ T cells in promoting HFD-induced inflammatory disease (**Extended Data Fig. 6e,f**).

**Extended Data Figure 6.**
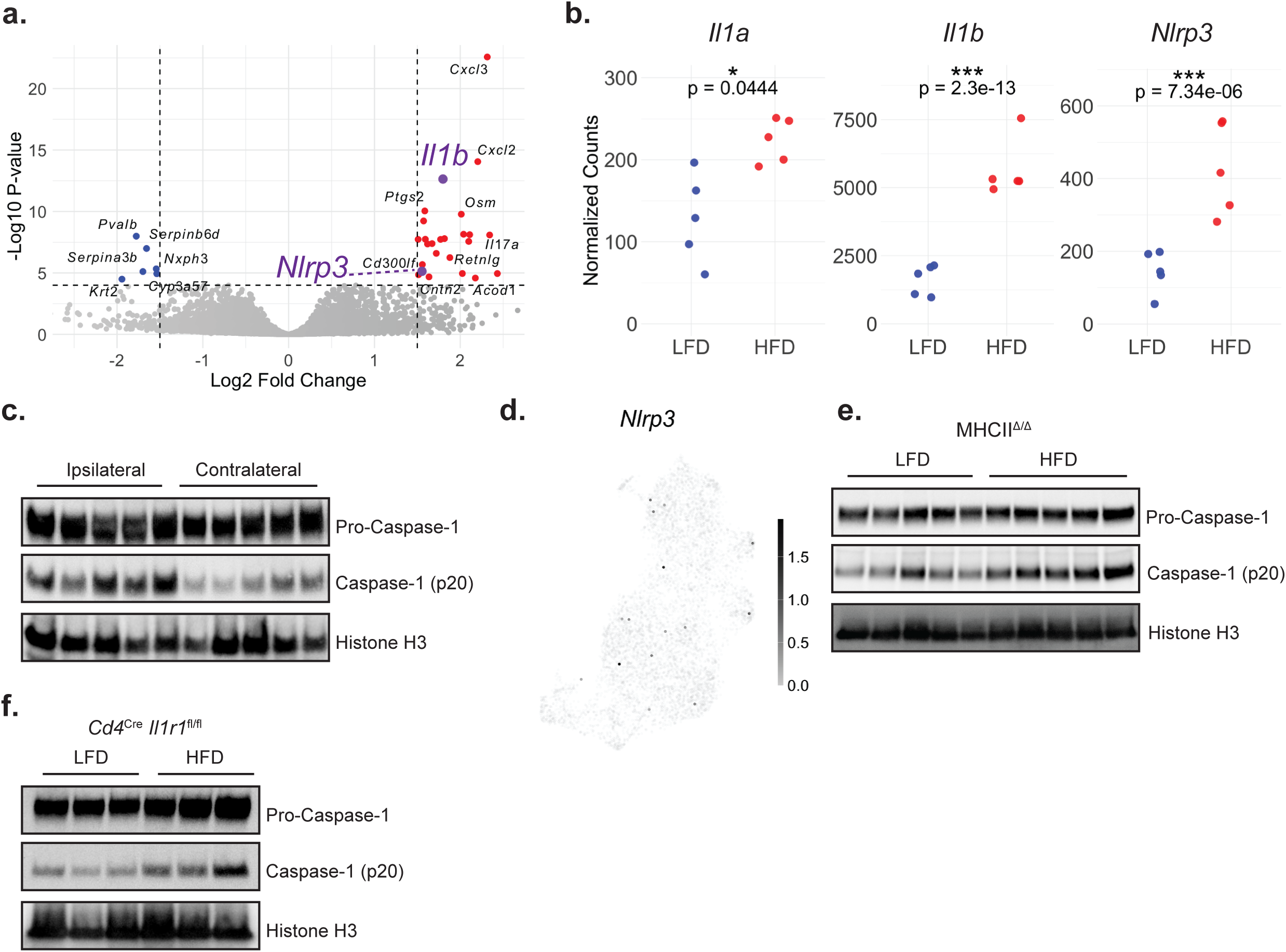
HFD-induced NLRP3 activation is extrinsic to CD4^+^ T cells, **a,** Volcano plot of bulk-RNA-seq of IMQ-treated ears of LFD or HFD-fed mice, **b,** Normalized counts of *II1a, II1b,* and *Nlrp3.* **c,** Western blot of Pro-Caspase-1, Capsase-1, and Histone H3 from Ipsilateral (left) and contralateral (right) skin draining lymph node lysates of WT mice fed on HFD. **d,** *Nlrp3* expression on UMAP plot from **Fig. 2I. e,f,** Western blot of Pro-Caspase-1, Capsase-1, and Histone H3 from skin draining lymph node lysates of MHCII^Δ/Δ^ mice **(e)** and *Cd4^Cre^ II1r1^fl/f l^*mice **(f)** treated with IMQ. n=5 **(a-c,e);** n=3 **(f).** DESeq2 Wald test was used to calculate *P* values and Rvalues were corrected for multiple corrections using the Benjamini-Hochberg (BH) method (a,b). *P < 0.05, **P < 0.01, ***P < 0.001.

To orthogonally validate the functional role of the NLRP3 inflammasome, we utilized MCC950, a well-established small molecule inhibitor of the NLRP3 inflammasome^48^. Six-week-old WT mice fed on HFD were topically treated with either vehicle or MCC950. MCC950 treatment effectively protected mice from HFD-induced exacerbation of psoriatic inflammation (**Fig. 3g,h**), as evidenced by a significant reduction in T_H_17 cells and neutrophils from the lesional skin (**Fig. 3i,j**). Consistently, topical MCC950 treatment also reduced HFD-induced NLRP3 activation in the lesional skin as monitored by diminished active caspase-1 (**Fig. 3k**). Altogether, these findings underscore the critical role of lesional NLRP3-dependent inflammasome activity in the rapid modulation of immune responses to obesogenic dietary cues.

### Transient HFD at disease onset has durable effects on disease recurrence

Psoriasis is a chronic inflammatory disease marked by onset, recovery, and repeated recurrence^49^. Emerging evidence suggests that inflammatory memory is retained in tissues after initial disease onset, which may facilitate disease recurrence in the same area after recovery^50–52^. To explore whether diet-on-disease onset (DODO) contributes to inflammatory disease trajectory beyond the onset phase to disease recurrence, we developed a psoriasis recurrence model (**Fig. 4a**). Mice were fed either LFD or HFD for 1 week, followed by daily IMQ challenge for 6 days while maintaining the LFD or HFD, respectively, mimicking the acute psoriasis model. Afterward, mice were given only LFD, and IMQ treatment was stopped. Following approximately 25 days of recovery, in which the ears appear normal by gross examination, ear thickness measurement, and histological examination, mice were subjected to a secondary IMQ challenge for 4 days (**Fig. 4a**). As expected, HFD-fed mice exhibited more severe inflammatory disease during the initial disease onset compared to LFD-fed mice (**Fig. 4b**). Notably, mice that had previously been on HFD (HFD^DODO^) continued to exhibit more rapid and pronounced recurrent inflammation – even though they were switched to LFD during both the recovery phase and the second flare. This was evidenced by a ∼2-fold increase in ear thickness, along with markedly increased T_H_17 cell infiltration and neutrophil accumulation in the skin (**Fig. 4b-e**). The T_conv_ cells from the lesional ears of HFD^DODO^ mice continued to express higher IL1R1 after the second flare (**Fig. 4f**). To determine whether HFD alone elicits persistent effects, mice were placed on HFD or LFD for one week without IMQ and then subjected to a recovery interval for two weeks on LFD. Following recovery, these mice were challenged with IMQ, and the mice previously fed HFD exhibited disease severity comparable to that of LFD-fed controls (**Extended Data Fig. 7a**), indicating that HFD in the absence of an inflammatory trigger is insufficient to produce a lasting impact. By contrast, transient HFD exposure during psoriasis onset profoundly augmented disease severity upon recurrence, underscoring the critical influence of dietary perturbations at disease onset in driving inflammatory memory and shaping long-term disease trajectories. These findings indicate that avoiding dietary triggers like fatty meals at disease onset might benefit those at risk or with established inflammatory conditions, potentially reducing the severity of future flares.

**Figure 4.**
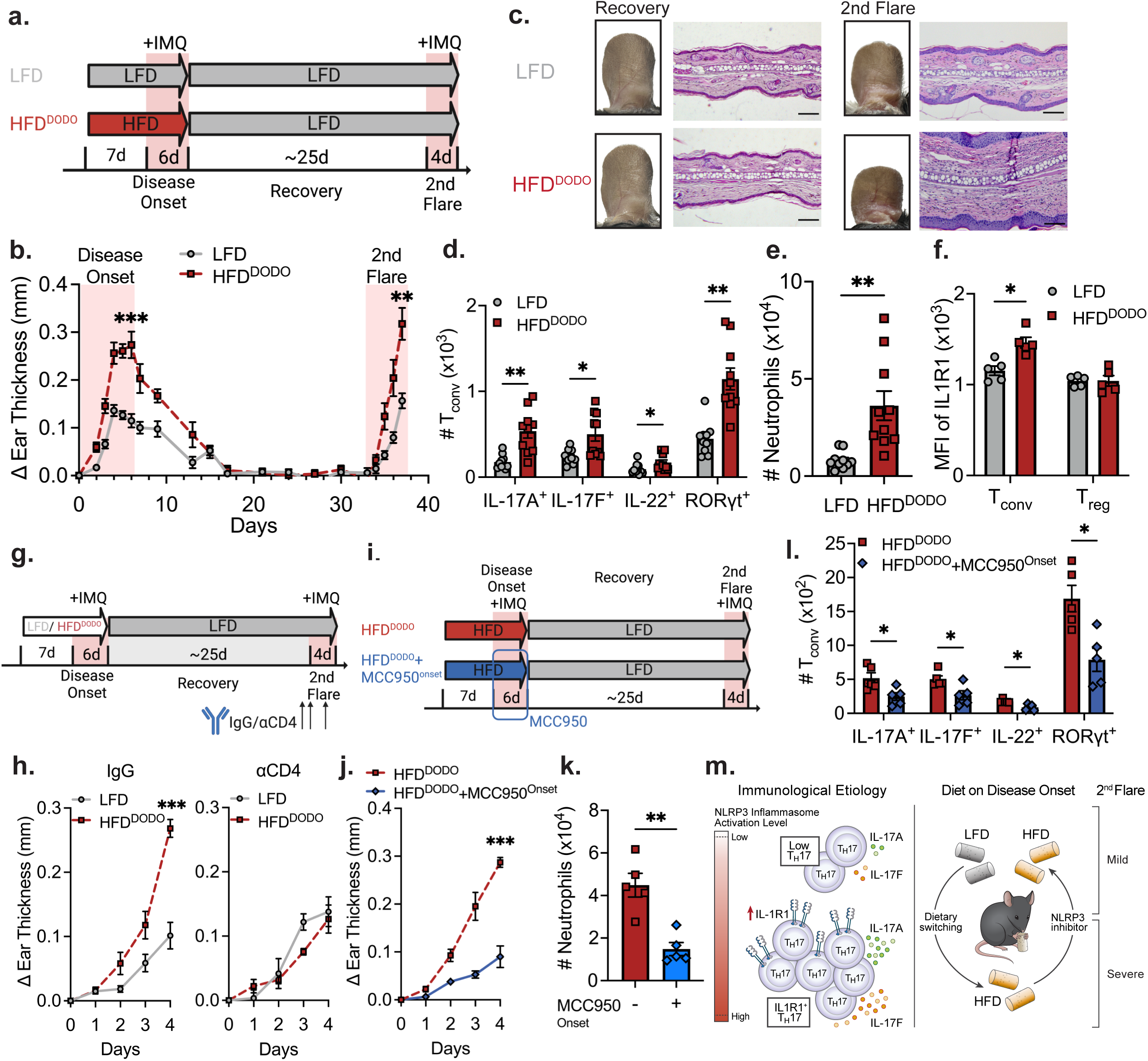
Diet-on-disease-onset (DODO) durably alters the severity of disease recurrence, **a**, Schematic of IMQ-induced psoriasis recurrence model, **b**. Change in ear thickness during the initial challenge, recovery phase, and 2^nd^ flare, **c**, Representative ear images and H&E ear histology of mice at end of recovery and after 2^nd^ flare, **d.e**. Total number of IL-17A^+^, IL-17F^+^. IL-22^+^, and ROR/t^+^ T_conv_ cells (**d**) and neutrophils (**e**) in the skin after 2^nd^ flare, **f**. MFI of IL1R1 in T_conv_ cells and Treg cells after 2^nd^ flare, **g**. Schematic of CD4^+^ depletion after recovery, **h**. Change in ear thickness during 2^nd^ flare from mice treated with IgG or anti-CD4 antibody after recovery. **I**, Schematic of MCC950 treatment during disease onset, **j-l**, Change in ear thickness (**j**), total number of neutrophils in the skin (**k**), and total number of IL-17A^+^. IL-17F^+^, IL-22^+^, and RORyt^+^ T_conv_ cells in the skin (**I**) after 2^nd^ flare in mice treated with vehicle or MCC950 during disease onset, **m**, Model of DODO as a critical modifier of disease onset and disease recurrence. Scale bars. 100 pm. n=5 (**b.f.h.j-l**); n=10 (**d.e**). Data are mean ± s.e.m. Welch’s West was used to calculate *P* values. *P* values were corrected for multiple corrections using Holm-SidSk method (**d.l**). Only peak values were tested in (**b.h.j**). *P < 0.05, **P < 0.01, ***P < 0.001.

**Extended Data Figure 7.**
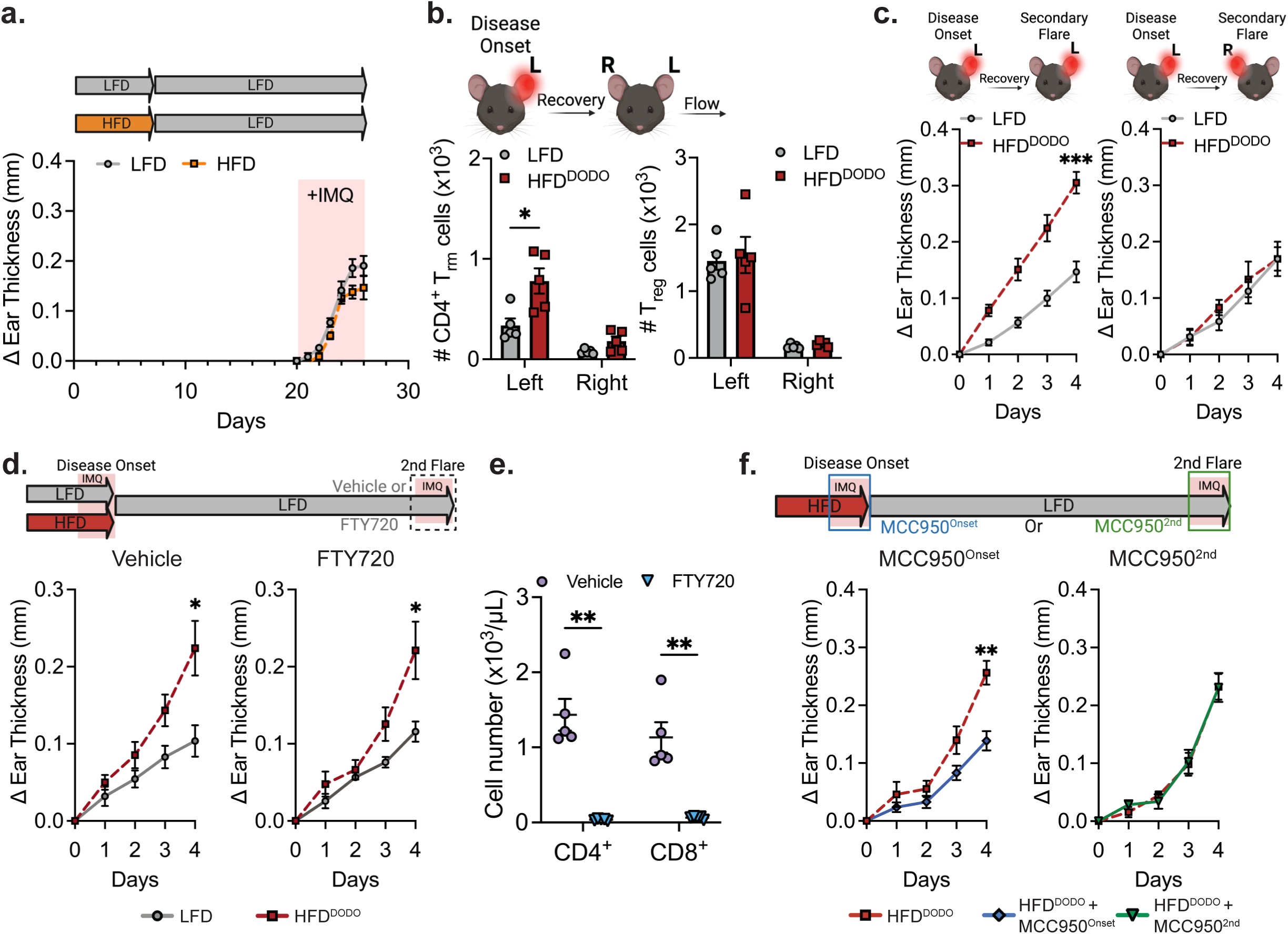
HFD exposure and therapeutic interventions shape psoriatic recurrence, **a**, Change in ear thickness of mice fed HFD or LFD for one week, followed by two weeks on LFD, and then challenged with IMQ while maintained on LFD. **b**, Total number of Trm (CD4^+^ FOXP3^−^ CD62L^low^ CCR7^low^ CD4high CD69^+^) and Treg cells from mouse ear skin after recovery, **c**, Change in left ear (previously challenged) or right ear (previously unchallenged) thickness upon rechallenge, **d**, Change in ear thickness during recurrence in mice treated with either vehicle or FTY720 on recurrence (2nd flare), **e**, Number of CD4^+^ and CD8^+^ T cells in peripheral blood from mice treated with either vehicle or FTY720. **f**, Change in ear thickness during recurrence in mice treated with MCC950 on disease onset (MCC950^Onset^) or MCC950 on recurrence (MCC950^2nd^). n=5, except in (**a**) where n=10 for mice previously fed on HFD for 1 week and (**d**) where n=4 for HFD-Vehicle. Data are mean ± s.e.m. Welch’s t-test was used to calculate P values. Only peak values were tested in (**a,c,d,f**). *P < 0.05, **P < 0.01, ***P < 0.001.

### DODO-induced tissue memory is driven by local NLRP3 activation at onset and the CD4^+^ compartment

To assess whether the durable effects induced by transient HFD feeding were mediated by inflammatory memory, we profiled the fully recovered ears (left) and unchallenged ears (right) before the second challenge. In the recovered ears of HFD^DODO^ mice, we observed an approximately 2.3-fold increase in resident memory CD4⁺ T (T_rm_) cells^53,54^, whereas the previously unchallenged ears showed no significant differences compared to LFD-fed controls (**Extended Data Fig. 7b**, **Extended Data Fig. 8**). Furthermore, when mice fully recovered from disease onset were subjected to a second flare, only the previously challenged (left) ears, but not the previously unchallenged (right) ears, exhibited a significant increase in ear thickness in HFD^DODO^ mice (**Extended Data Fig. 7c**), suggesting that the durable immunomodulatory effects of DODO are localized and lesioned tissue-specific. In addition, blocking lymphocyte egress with FTY720^55,56^ during the second flare did not prevent HFD^DODO^-induced exaggerated inflammation, further suggesting a tissue-resident immune response (**Extended Data Fig. 7d,e**).

To determine whether the molecular events that potentiate more severe inflammatory disease upon transient HFD exposure at disease onset – specifically, NLRP3 activation driving CD4^+^ T cell inflammation – also underlie disease recurrence, we targeted CD4^+^ T cells in mice that had recovered from their initial flare (**Fig. 4g**). Intraperitoneal administration of anti-CD4 or isotype control (IgG) antibodies revealed that whereas HFD^DODO^ mice treated with IgG displayed a marked exacerbation of disease severity upon re-challenge, anti-CD4 treatment abolished this difference between HFD^DODO^ and LFD-fed mice (**Fig. 4h**). These results establish CD4^+^ T cells as essential mediators of the durable inflammatory memory induced by transient HFD feeding. Further, topical administration of MCC950 during disease onset significantly attenuated the heightened inflammation observed in HFD^DODO^ mice during recurrence, compared with HFD^DODO^ mice treated topically with PBS (**Fig. 4i–l**). Notably, identical topical MCC950 treatment applied solely during the recurrence phase failed to confer comparable protection (**Extended Data Fig. 7f**), underscoring a discrete therapeutic window in which local inhibition of skin NLRP3 activation can prevent diet-induced priming of pathogenic T_H_17 responses and forestall more severe disease upon relapse. In addition to avoiding certain dietary triggers such as fatty meals, we speculate that topical NLRP3 inhibition at disease onset might represent a viable strategy to prevent a heightened inflammatory response during future flares. Overall, our findings reveal that transient exposure to an obesogenic diet at disease onset – rather than chronic obesity per se – reprograms the immune milieu to promote durable inflammatory memory and exacerbate disease severity, suggesting a shift in our understanding of obesity-associated immune dysfunction (**Fig. 4m**).

**Extended Data Figure 8.**
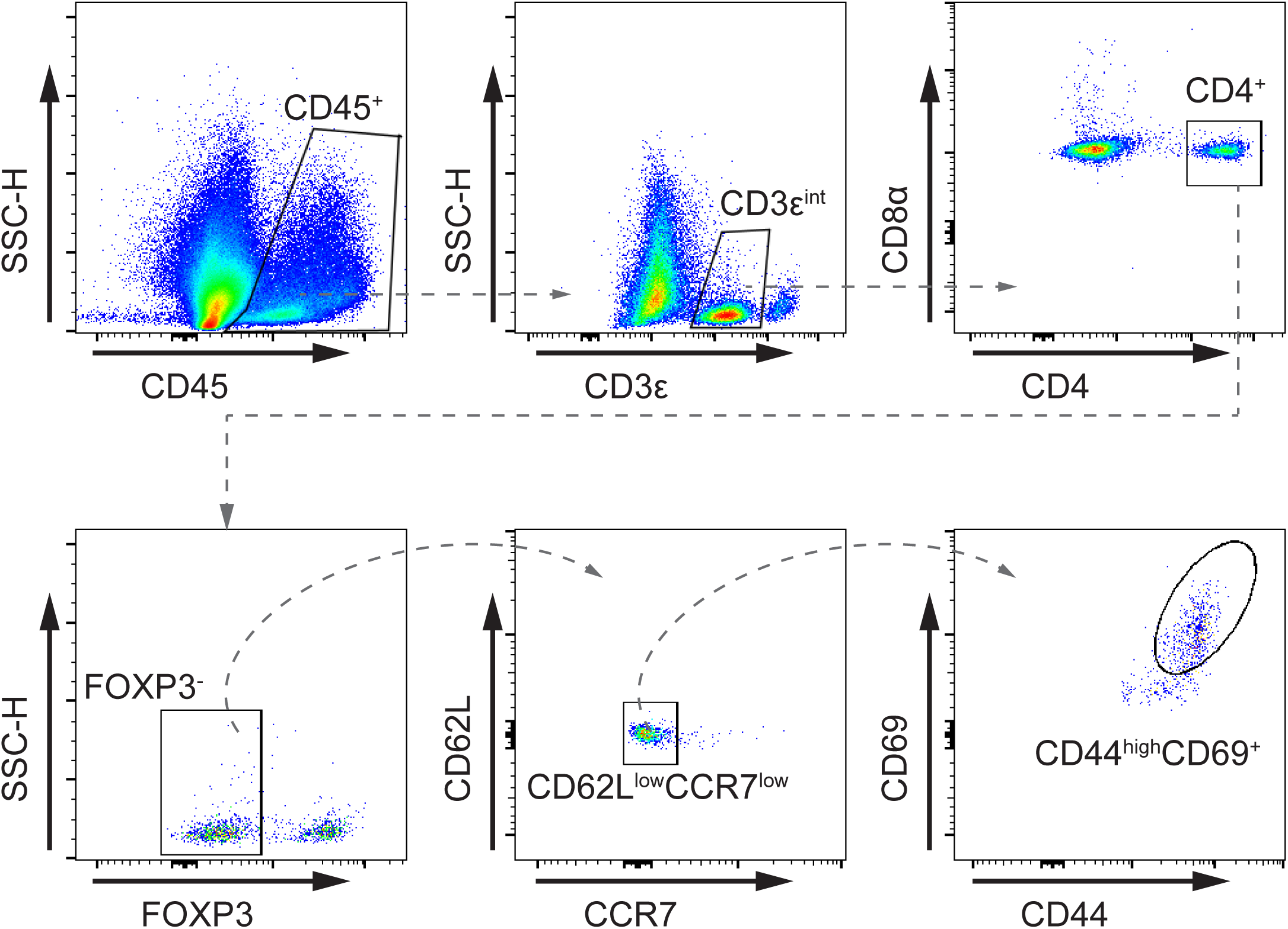
Gating flow cytometry gating strategy to quantify T_rm_cells. Specifically, resident memory CD4^+^ T cells (T_rm_ are defined as CD45^+^ CD3ε^Int^ CD4^+^ FOXP3^−^ CCR7^low^ CD62L^low^ CD44^high^ CD69^+^.

## Material and Methods

### Mice

Mouse studies were conducted at the University of California, San Francisco (UCSF), USA. Mice were housed in specific pathogen-free facilities. When required, mice were purchased from The Jackson Laboratory. By default, mice fed at UCSF were given irradiated PicoLab Rodent Diet 20 5053. For experiments requiring the use of special diets, diets were irradiated and purchased from Research Diets. The HFD used was 60 kcal% fat (D12492), the control LFD was 10 kcal% fat (D12450J). Mice designated as obese in this study were fed HFD for at least 10 weeks, starting at an age of 6–7 weeks, and weighed at least 38 g (as an average within a cage). For short-term HFD feeding, 6-week-old mice were given HFD for 7 days before the imiquimod (IMQ) challenge unless otherwise indicated. For HFD^in^ feeding, mice were placed on HFD starting two days before IMQ challenge and remained on it until two days after initiation. For HFD^out^ feeding, mice were maintained on LFD except for two distinct periods when they were switched to HFD: first, from seven to four days before IMQ initiation, and later, from four to six days after initiation during disease progression. For HFD^DODO^ feeding, mice were placed on HFD throughout the entire disease onset phase (spanning seven days before to six days after IMQ initiation) and were then returned to LFD for the recovery and rechallenge phases. C57BL/6J (#000664), B6.SJL-Ptprca Pepcb/BoyJ (002014)^57^, *Rag1*^−/−^ (#002216)^58^, *Tcra*^−/−^ (#002116)^59^, B6.129S2-H2dlAb1-Ea/J (MHC II^Δ/Δ^) (#003584)^60^, *B2m*^−/−^ (#002087)^61^, *Tcrd*^−/−^ (#002120)^62^, *Cd4^Cre^* (#022071)^63^, and *Nlrp3*^−/−^ (#021302)^64^ mice were purchased from The Jackson Laboratory. *Il1r1^fllfl^* ^65^ mice were kindly provided by Scharschmidt Lab at UCSF. All mice used for the studies were male.

All procedures involving animals were performed in accordance with protocols approved by the Institutional Animal Care and Use Committee (IACUC) of UCSF.

### IMQ-induced psoriasis mouse model

IMQ-induced psoriasis was generated in mice at various ages fed with either LFD or HFD. In brief, mice were anesthetized using isoflurane, and 15 mg cream containing 5% IMQ (TARO) was topically applied to the dorsal and ventral sides of the left mouse ear daily for 6 days^23^. Ear thickness was measured daily with a digital micrometer (Mitutoyo). 24 h after the last application, mice were sacrificed, and the ears were collected for flow cytometry, RNA extraction, histological analysis, and other downstream applications. For rechallenge experiments, after the last day of IMQ treatment during the initial challenge, mice were allowed to recover. Ear thickness was measured weekly to evaluate recovery. After 25-30 days, mice were fully recovered from previous inflammation and were subjected to 15 mg cream containing 5% IMQ treatment for 4 consecutive days. To inhibit NLRP3 activation, MCC950 sodium (#S7809, Selleckchem) was prepared in 50 mg mL^−1^ in water and 75 μg was given topically to each mouse daily in sync with application of IMQ.

### Single-cell suspension of ear-infiltrating immune cells

Dissected mouse ear skin was separated into dorsal and ventral halves and incubated in 1 mL of digestion buffer. Mouse skins were incubated in 1 mL digestion buffer (RPMI-1640 media with 2% fetal calf serum and 1x Penicillin-Streptomycin-Glutamine solution (#10378016, Roche) containing 0.2 mg mL^−1^ Liberase (#05401020001, Roche), 0.2 mg mL^−1^ DNase I (#DN25-100MG, Sigma), and 0.5 mg mL^−1^ Hyaluronidase (#LS002592, Worthington). After 30 min incubation, skin samples were minced into fine pieces (1–2 mm^3^) and incubated for another 30 min. The suspension was then passed through a 100 μm mesh to remove undigested clumps and debris. The flow-through was centrifuged at 500 RCF for 5 min. The pellet containing the stromal vascular fraction was washed once in 10 mL RPMI, and the resultant isolated cells were prepared for FACS analysis directly or first stimulated with PMA and ionomycin in the presence of brefeldin A (Cell Activation Cocktail; Biolegend) for 1.5 h at 37 °C for subsequent intracellular cytokine staining and then prepared for FACS analysis.

### *In vivo* antibody treatments

The following antibodies were used: anti-mouse CD4 (BioXCell, GK1.5) and isotype control for anti-mouse CD4 (Rat IgG2b, BioXCell, LTF-2). The antibodies were injected intraperitoneally two days before, on the day of, and two days after the initiation of the IMQ-induced psoriasis challenge at a dose of 0.1 mg per mouse. For the rechallenge experiment, the anti-CD4 or isotype control antibodies were injected intraperitoneally on days 29, 32, and 35 following the initial IMQ challenge using the same dosage. Injection volumes never exceeded 150 μl.

### Adoptive transfer of CD4^+^ T cells

CD4^+^ T cells were isolated from the spleens of WT mice using Mouse CD4^+^ T Cell Isolation Kit (Stemcell) and 100,000 cells were transferred intravenously into *Rag1*^−/−^ mice. Following an 8-week reconstitution period, mice were fed LFD or HFD for 1 week. Mice were then subjected to IMQ treatment for 6 days to induce psoriatic-like skin inflammation. Disease severity was assessed via ear thickness measurements, and immune responses were analyzed by flow cytometry.

### Flow cytometry

Fc Block (2.4G2, #553141, BD) was diluted in FACS buffer and added to each sample 5 minutes prior to performing surface stain. eBioscience Fixable Viability Dye eFluor780 was used to distinguish live from dying or dead cells. Cell surfaces were stained for 30 minutes at 4 °C. For intracellular staining, cells were treated with fixation and permeabilization reagents from eBioscience and labeled with appropriate antibodies before being analyzed. Data was analyzed using Aurora Cytek and FlowJo software (FlowJo LLC). Cells were sorted with Cytek Aurora™ CS System. The following antibodies were used: CD45.2 (104, #109839, Biolegend), CD90.2 (30-H12, #740205, BD), CD197 (4B12, #120120, Biolegend), CD11c (N418, #117322, Biolegend), CD44 (IM7, #103037, Biolegend), TCRγδ (GL3, #563993, BD), CD62L (MEL-14, #20-0621-U025, TONBO), CD69 (H1.2F3, #104506, Biolegend), Ly6G (1A8, #127672, Biolegend), CD3ε (145-2C11, #100349, Biolegend), CD4 (RM4-5, #100528, Biolegend), CD8α (53-6.7, #100744, Biolegend), Ki-67 (SolA15, 367-5698-82, Invitrogen), RORγt (Q31-378, #562684, BD), FOXP3 (FJK-16s, #46-5773-82, Invitrogen), IL-17A (TC11-18H10.1, #506914, Biolegend), IL-17F (9D3.1C8, #517008, Biolegend), CD121a (35F5, #564387, BD), IFNγ (XMG1.2, #505806, Biolegend), IL-22 (Poly5164, #516409, Biolegend).

### Tissue RNA isolation and bulk-RNA sequencing

Total RNA was extracted from the whole mouse ear skin before and post IMQ challenge using a RNeasy Fibrous Tissue Kit according to the manufacturer’s protocol (#74704, Qiagen). RNA quantity and quality was determined by NanoDrop spectrophotometers and Qubit 4 fluorometer. 10 μg RNA from each sample was sent to Genewiz for library preparation and sequencing services (RNA-seq). Read alignment and junction finding was accomplished using STAR^66^ utilizing UCSC mm10 as the reference sequence. Differential gene analysis and data visualization was performed in R using DESeq2^67^ package.

### scRNA-Seq of T cells isolated from lesional skin

CD4^+^ T cells from mouse ear skin were FACS-sorted and then mRNA isolation, cDNA synthesis, and library preparation were performed based on the protocol provided by PIPseq™ T2 3ʹ Single Cell RNA Kit v4.0PLUS (Fluent BioSciences). In brief, cells were sorted by FACS (sorting on live CD45.2^+^ CD3ε^+^ CD4^+^ cells on a Cytek Aurora™ CS System) in Cell Suspension Buffer (provided in the kit) with 1 U μL^−1^ RNase inhibitor. 4 μL cell suspension (5000 cells max) with 20 units of RNase inhibitor was added to each PIP tube reaction, followed by vortexing to generate single bead to cell emulsion. We performed 16 cycles of PCR for cDNA amplification after cDNA synthesis, and 25% of each cDNA sample was carried into transcriptome library preparation. PIPseq T2 3ʹ Single Cell gene expression libraries were subject to standard Illumina paired-end sequencing targeting 20,000 reads per input cell using NovaSeq X.

Raw fastq files were mapped to the mouse transcriptome (UCSC mm10) using PIPseeker^68^ (Fluent BioSciences, version 3.2) pipeline that performs barcode whitelisting, alignment with STAR, and molecular index correction, and generates barcode-feature counts matrices. After alignment and sensitivity calling (sensitivity_5 was chosen for all three samples), 2103 cells were recovered from the LFD condition and 5232 cells were recovered from the HFD condition – two T2 Pipseq reactions were used, labeled HFD1 and HFD2. After QC filtering (for doublets and non-CD4^+^ T cell contamination), 1769 CD4^+^ T cells remained in the LFD condition, and 4450 CD4^+^ T cells remained in the HFD condition, with over 1,500 median genes per cell. Seurat^69^ (version 5.1.0) was used for downstream analysis. The filtered expression matrices for LFD, HFD1, and HFD2 samples were loaded and processed to create Seurat objects for each condition. The HFD1 and HFD2 datasets were merged into a single HFD Seurat object, which was subsequently merged with the LFD dataset. The combined dataset was then normalized using the NormalizeData function with default settings. Variable features were identified using the “vst” method with 2,000 features. Data scaling was performed using ScaleData, followed by principal component analysis (PCA) using the top 2,000 variable features. To correct for batch effects between the different conditions, Harmony^70^ (version 1.2.0) was applied on the PCA embeddings, using the condition labels (LFD, HFD1, HFD2) as a covariate. Batch-corrected embeddings were used for downstream analyses, including clustering and visualization. Clustering was performed by constructing a shared nearest neighbor (SNN) graph using the FindNeighbors function on the Harmony-corrected embeddings, followed by clustering using the FindClusters function with a resolution of 0.5. Uniform Manifold Approximation and Projection (UMAP) was used for visualization of the clusters, based on the Harmony embeddings. For downstream analysis, clusters identified as potential contaminations (clusters 9 and 10, likely monocytes and stromal cells) were removed, and clustering was redone. Differential expression analysis was conducted using the FindAllMarkers function with the following parameters: only positive markers (only.pos = TRUE), minimum fraction of cells expressing a gene (min.pct = 0.25), and a log fold change threshold of 0.25. The top five markers for each cluster were identified and visualized using a dot plot (DotPlot). To assess T_H_17 pathogenicity, we calculated a T_H_17 pathogenicity score based on published gene signatures associated with pathogenic T_H_17 cells^8,39^. Gene sets from both studies, comprising key regulators and effector genes that define pathogenic T_H_17 cells, were curated and used to evaluate the pathogenic potential of T_H_17 cells in our dataset. We first identified the genes present in our single-cell RNA-seq dataset that overlapped with the published pathogenic T_H_17 gene sets. Using the AddModuleScore function in Seurat, we calculated the T_H_17 pathogenicity score for each cell in the T_H_17 cluster.

### Histology of lesional ear skin

Ear skin was harvested and submerged in 4% paraformaldehyde overnight. Tissues were then embedded, sectioned, and stained with hematoxylin and eosin (H&E) by the Histology and Light Microscopy Core at Gladstone Institutes. Images were taken using a Zeiss microscope. The scale bar was added by converting pixels to microns using ImageJ software.

### Western blots of caspase-1

For western blotting, homogenized tissues or whole-cell lysates, samples were lysed in RIPA buffer containing protease inhibitor cocktail (Roche). Cell lysates were centrifuged at 14,000 x RCF for 15 min, and supernatants were prepared in 4X LDS Sample Buffer (Invitrogen), separated by SDS-PAGE, and transferred to Immobilon 0.45μm membranes (Millipore). All primary antibody incubations were performed overnight at 4 °C and secondary antibody incubations were performed at room temperature for 2h. All primary antibodies were diluted at 1:1000 in TTBS supplemented with 3% BSA. Secondary antibody incubations were performed at 1:10,000 dilution in 1:1000 in TTBS supplemented with 5% nonfat milk. The following primary antibodies were used for the study: anti-Caspase-1 (p20) mAb (#AG-20B-0042-C100, AdipoGen) and anti-Histone H3 pAb (#9715, Cell Signaling).

### FTY720 treatments

Mice received daily intraperitoneal injections of 100 µl/mouse of FTY720 (1 mg/kg) or water (vehicle), starting one day before the IMQ challenge and continuing daily through its duration. This treatment was applied during either the initial disease onset or the rechallenge phase.

### Statistical analyses

Statistical analyses were performed with Prism 10 (GraphPad). *P* values were calculated using Welch’s *t*-test unless otherwise noted. When applicable, mice were randomly assigned to treatment or control groups. Experimental data was not excluded from the statistical analyses, except for exclusions due to technical errors or loss of confidence in appropriate controls. Measurements were taken from distinct biological samples, except for ear thickness, which was measured longitudinally in the same mice at multiple time points. The investigators were not blinded in the studies unless otherwise stated. No estimate of variance was made between each group. All *in vivo* experiments were conducted with at least two independent cohorts unless otherwise specified. Unless otherwise noted, data from one instance of the repeated experiments are represented in this manuscript. Bulk and single-cell RNA-seq experiments were conducted once using multiple biological samples per group (as indicated in figure legends).

## Acknowledgments

We thank A. Garg, R. Ricardo-Gonzalez, P. Dumesic, and J. Cyster for advice and critical review of the manuscript, S. Pyle for scientific graphic illustration, and Life Science Editors for editing services. Z.J. was supported by the UCSF Biomedical Sciences graduate program, UCSF Diabetes Center Klein Fellowship, and George Coleman Discovery Fellowship. S.P.B. holds a Career Award for Medical Scientists from the Burroughs Wellcome Fund and has been supported by U.S. National Institutes of Health (NIH) grants R38 HL143581 and K38 HL154202 as well as support from the Larry L. Hillblom Foundation, Parker B. Francis Foundation, the Baszucki Lymphoma Therapeutic Initiative, the Parker Institute for Cancer Immunotherapy, the Benioff Center for Microbiome Medicine, the UCSF Nutrition Obesity Research Center, and the American Partnership for Eosinophilic Disorders. We thank Dr. Tiffany Scharschmidt for sharing *Il1r1^fl/fl^* mice.

## Contributions

Z.J. and S.P.B. designed and conceived the research. S.P.B., A.J.C., and J.M.G. supervised the research. Z.J., J.M.T., C.T., S.G.G., A.Y., I.S., J.W., S.S.C., P.D., V.J., B.D., and J.T. performed experiments. Z.J., H.Y., N.F.K, T.C., A.J.C, and S.P.B. analyzed data. Z.J. and S.P.B. wrote the manuscript.

## Data availability

The raw bulk RNA-seq data generated in this study have been deposited in the NCBI Sequence Read Archive (SRA) under the BioProject accession number PRJNA1212208. Additionally, the raw and processed scRNA-seq data generated in this study have been deposited in the Gene Expression Omnibus (GEO) repository under the accession number GSE287431. Other relevant data are available from the corresponding author upon reasonable request.

## Code Availability

Code used in this manuscript can be provided by reasonable request to the corresponding author.

